# SIMBA-GNN: Simulation-augmented Microbiome Abundance Graph Neural Network

**DOI:** 10.1101/2025.05.27.656377

**Authors:** Mohammad Parsa, Javad Aminian-Dehkordi, Mohammad R.K Mofrad

**Affiliations:** Department of Bioengineering, University of California, Berkeley, California, US

**Author notes:** Equal contribution (Mohammad Parsa, Javad Aminian-Dehkordi). Correspondence to: Mohammad Mofrad < >.

**Keywords:** Graph neural network, Heterogeneous graph transformer, Gut microbiome, Cross-feeding, Metabolic networks, Edge-aware attention

## Abstract

Understanding gut microbiome dynamics gut requires deciphering complex, metabolically driven interactions beyond taxonomic profiles. We present `SIMBA`, a novel framework that integrates mechanistic metabolic simulations with a graph neural network (GNN) to predict microbial abundances and uncover cross-feeding relationships. By simulating pairwise interactions among gut microbes using metabolic networks, we generate biologically grounded graphs that capture metabolite cross-feeding and functional relationships. Our custom GNN, enhanced with edge-aware attention, is trained through a multi-stage pipeline combining self-supervised learning, simulation-based pretraining, and fine-tuning on real microbial abundance data. `SIMBA` achieves state-of-the-art performance (Spearman ρ = 0.85) and enables interpretable insights into keystone taxa and metabolic bottlenecks. This work demonstrates the power of combining metabolic networks with deep learning for precision microbiome analysis.

## 1. Introduction

The human gut microbiome is a dynamic ecosystem that modulates host physiology and has been linked to various diseases, from metabolic disorders to inflammatory and neurodegenerative conditions. These associations stem not from individual microbes in isolation, but from complex, context-dependent interactions within the microbial community. Deciphering these interactions is key to uncovering the mechanisms underlying microbiome-associated health outcomes and guiding the development of targeted therapies (Dong & Mayer, 2024; Cho et al., 2024).

Conventional approaches to microbiome analysis often rely on static representations, such as taxonomic abundance profiles or co-occurrence networks (Wang et al., 2021). While useful, these methods are limited in capturing the dynamic, nonlinear interactions, such as metabolic cross-feeding and competition, that shape microbial community behavior (Quinn-Bohmann et al., 2025; Aminian-Dehkordi et al., 2022). Recent advances in machine learning, particularly graph neural networks (GNNs), offer a powerful alternative for learning the structured, relational data inherent to microbial ecosystems.

GNNs have shown remarkable success in various biological applications, including modeling protein-protein interactions, disease classifications from microbial abundance profiles (Sun & Song, 2024; Zeng et al., 2024), and drug susceptibility prediction in microbiomes (Rehman et al., 2024). Additionally, studies have shown the utility of GNNs in analyzing gut microbiome metaomic data, providing insights into disease phenotypes (Irwin et al., 2024). Their ability to learn directly from graph-structured data makes them especially well-suited for analyzing microbial interaction networks, where nodes represent species and edges encode biological relationships.

However, current GNN-based approaches in microbiome research face significant limitations. Many depend on predefined or co-occurrence-based graphs that may not reflect true biological interactions, lack integration with mechanistic models, and exhibit limited generalizability beyond the training context. Moreover, progress is constrained by a shortage of high-quality, labeled datasets that capture microbial interactions under physiologically relevant conditions (Tian et al., 2023).

To address these challenges, we introduce `SIMBA-GNN`, a novel framework that integrates mechanistic metabolic simulations of microbial interactions with a GNN-based learning algorithm to model and predict the behavior of gut microbial communities. Specifically, we simulate pair-wise interactions between microbial species under realistic dietary constraints to generate mechanistically grounded datasets that capture emergent ecological phenomena such as cross-feeding patterns. Each simulation yields a graph where nodes represent microbial species, metabolic pathways, metabolites and edges encode the direction, microbe similarity, and the probability of the metabolites being produced and consumed. These graphs form a basis of our training data for a custom-designed GNN, which is trained to predict community-level properties, including microbial abundances and potential keystone interactions.

To ground this approach in real-world microbiome contexts, we use a large dataset of fecal metagenomes from individuals on a high-fiber diet, and simulate all pairwise interactions among microbes using metabolic networks. These simulations generate a rich interaction dataset under diet-constrained conditions, enabling us to construct labeled graphs capturing microbial cross-feeding dynamics. Our GNN is then trained using these graphs and evaluated for its ability to predict observed microbial community compositions and highlight critical cross-feeding relationships.

Our contributions are as follows:

- A novel computational pipeline that generates a large-scale dynamic dataset of pairwise microbial cross-feeding interactions using metabolic networks constrained by realistic dietary inputs.
- A custom GNN architecture tailored to learn from simulation-derived microbial interactions for the prediction of microbial abundances.
- Comprehensive evaluation of the model’s predictive performance on microbial abundances and interpretability, demonstrating the potential of integrating metabolic networks with advanced deep learning for understanding complex microbial ecosystems.

This study opens new avenues for personalized microbiome modeling and precision therapeutics by providing a scalable, mechanistic approach to understanding the human gut microbiome ecosystem.

## 2. Methods

### 2.1. Microbial abundance data and selection of GEMs

Microbial abundance profiles were obtained from 186 individuals participating in a high-fiber dietary intervention study (Diener et al., 2020). This experimentally microbial abundance data served as the target variable for our GNN model. All identifiable microbial taxa from the cohort’s microbiome profiles were extracted. For each taxon, corresponding genome-scale metabolic models (GEMs) were retrieved from the AGORA database (version 1.03) (Heinken et al., 2020), a manually curated repository of semi-automatically reconstructed GEMs for common hu-man gut microbes. This resulted in a collection of 76 unique GEMs representing key members of the cohort’s gut microbial community. Each GEM encodes a stoichiometric reconstruction of its corresponding organism’s metabolic network, which enables simulation of metabolic fluxes. The GEMs were processed using COBRApy.

### 2.2. Simulation of pairwise microbial interactions

To systematically investigate potential cross-feeding interactions, we performed pairwise co-culture simulations for all 2,850 unique combinations of the selected GEMs.

#### 2.2.1. Model merging and shared environment

For each pair of microbes, their corresponding GEMs were combined into a single multi-compartment model. Metabolites and reactions from each organism were uniquely labeled to avoid conflicts, and a shared *pool* compartment was introduced to facilitate metabolite exchange. For each overlapping exchange reaction present in both original models, a corresponding metabolite was created in the pool compartment. Transport reactions were added to allow each microbe to secrete into or uptake from this shared metabolite pool, allowing simulation of potential cross-feeding interactions.

#### 2.2.2. Dietary constraints

The simulations were constrained to mimic a high-fiber diet. An averaged high-fiber diet profile, based on experimental data (Diener et al., 2020), was used to define the input metabolic fluxes into the shared lumen compartment. Simulations were performed under anaerobic conditions, reflecting the dominant environment of the human colon. Non-dietary exchange reactions were initially closed for uptake, allowing only secretion into the lumen, unless a metabolite was part of the defined medium. Metabolic byproducts were allowed to exit the system to simulate realistic environmental turnover (upper bounds of exchange reactions typically set to 1000).

#### 2.2.3. Objective functions and flux sampling

For each pairwise simulation, the objective was set to maximize the joint biomass production of the pair. To ensure both microbes could grow, the individual biomass reaction of each microbe was constrained to be at least 10% of its optimal growth rate when simulated in monoculture under the same dietary conditions. Flux sampling was then performed on the combined model using the ’optgp’ method (Megchelenbrink et al., 2014), generating 10,000 flux distributions with a thinning factor of 100. This process explores the feasible flux space under the given constraints and objective. The Geweke diagnostic test was applied to the sampled flux distributions for each reaction to ensure the convergence of the sampling process.

#### 2.2.4. Identification of cross-feeding metabolites

Cross-feeding metabolites were identified from the sampled flux distributions of the pairwise models. A metabolite was classified as cross-fed if it was secreted by one microbe and concurrently taken up by the other *via* the shared lumen compartment. To quantify this interaction, Spearman correlation analysis was performed on the exchange fluxes of the paired microbes for each shared metabolite. A strong negative correlation (*e*.*g*., threshold ¡ −0.5) was interpreted as indicative of potential cross-feeding. For each event, the directionality and frequency (*i*.*e*., number of samples showing the exchange) were recorded. Only metabolites with exchange fluxes exceeding a certain threshold and not present in the initial dietary input were retained. The presence probability for each identified cross-fed metabolite, quantified as the proportion of exchange events observed across 10,000 flux samples, was computed and incorporated as an edge feature for the next step.

#### 2.2.5. Pathway activity scoring

To evaluate functional metabolic contributions, we quantified pathway-level activity in each microbe based on flux distributions from pairwise simulations. Reactions were mapped to metabolic pathways using annotations from the Virtual Metabolic Human (VMH) and KEGG databases. Pathway activity scores, hereafter called pathway fingerprints, were calculated by aggregating the fluxes of constituent reactions, enabling pathway-level comparisons across microbial interactions (Figures S1 and S2).

### 2.3. Graph Neural Network Architecture

#### 2.3.1. Microbial Interaction Graph Construction

We constructed microbial interaction graphs based on 2,850 pairwise metabolic simulations. Each simulation yielded metabolite fingerprints, capturing the probability of individual metabolites being produced or consumed between microbial pairs as well as pathway fingerprints, summarizing pathway-level activities for each microbe.

Our heterogeneous graphs encoded three node types—microbes, metabolites, and pathways—and multiple biologically meaningful edge types. Microbe–metabolite edges encode directional interactions based on metabolite exchange probabilities inferred from simulations, effectively modeling ecological interactions such as cross-feeding. Microbe–pathway edges represent functional associations, linking microbes to pathways with nonzero activity. In addition, microbe–microbe edges were added based on pathway profile similarity, using cosine similarity between log-transformed pathway activity vectors. Only edges with similarity score above 0.85 were retained to reduce spurious connections and maintain graph sparsity. (Figure 2b)

**Figure 1.**
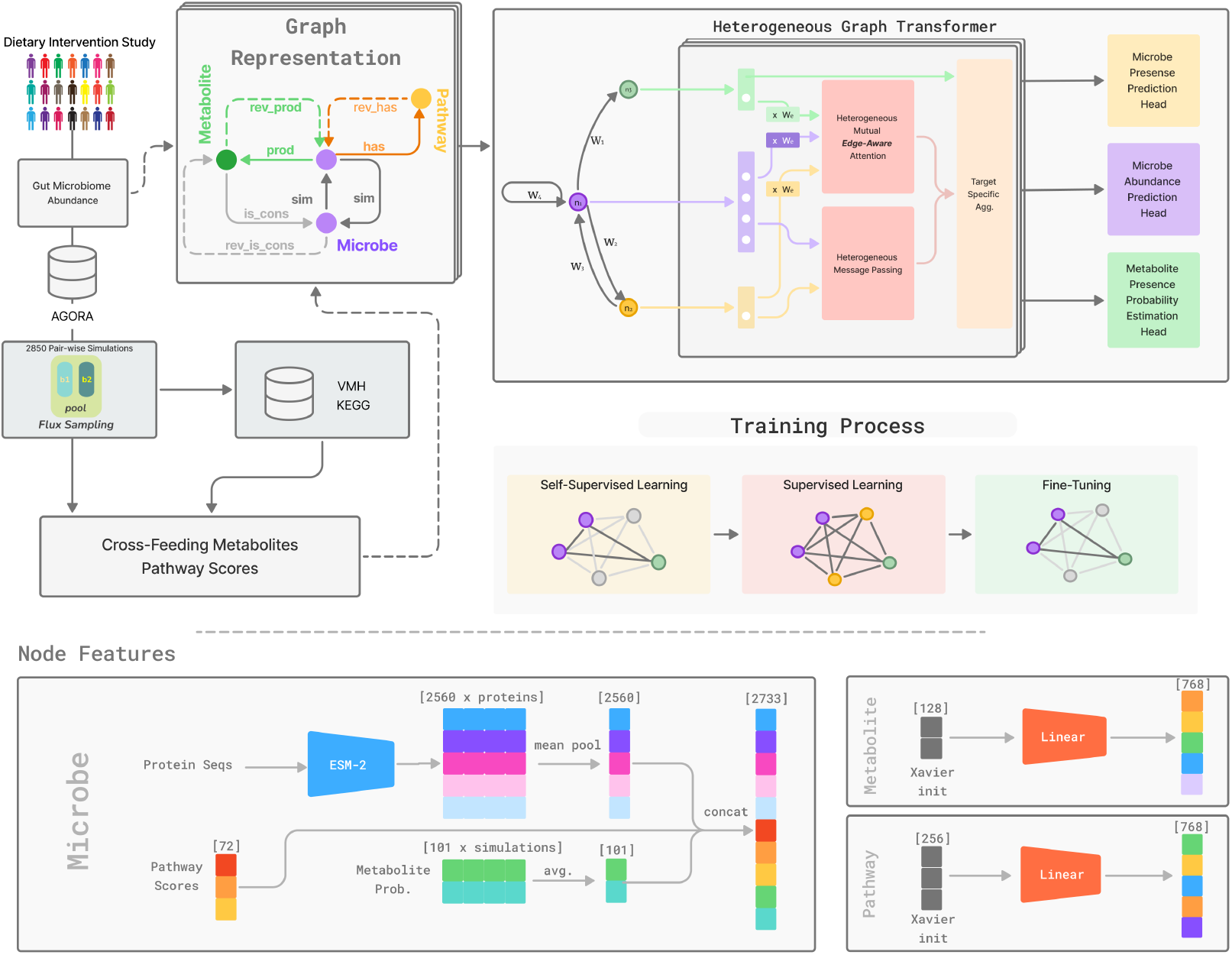
Overview of `SIMBA` **Left – Data sources**. A dietary intervention cohort of 186 individuals provides per-sample microbial relative abundances, which are mapped to their corresponding metabolic networks. 2850 pairwise simulations using flux sampling yield cross-feeding metabolites probabilities (used as metabolite fingerprints) and pathway activity scores (used as pathway fingerprints) between microbial pairs. All three signals are integrated into a unified dataset capturing cross-feeding interactions and metabolic pathways. **Middle-left – Graph construction**. We build a heterogeneous graph with three node types—microbe, metabolite, and pathway—connected by seven directed edge types: (i) `has`/`rev has` (microbe–pathway membership), (ii) `sim` (microbe–microbe cosine similarity *>* 0.85 based on pathway fingerprints), and (iii) `prod`, `cons` with their reverse edges, whose weights *w*_*e*_ = log_1+_ |flux| encode interaction strength. **Middle-right – Heterogeneous graph transformer**. Node features are first projected to 768-d and passed through three layers of our custom *edge-aware* HGT. Attention scores are modulated by scalar edge weight *w*_*e*_.The model outputs feed three task-specific heads: a BCE presence classifier, a per-sample softmax abundance regressor, and a BCE metabolite-probability estimator. **Bottom – Training schedule**. The network is optimized in three stages: (i) *Self-supervised* GraphCL to initialize embeddings, (ii) *Supervised pretraining* on simulated graphs with BCE, Tweedie (*p*=1.5) and metabolite BCE losses, and (iii) *Fine-tuning* on experimental graphs using only BCE and Tweedie losses (ranking loss tested but not retained). Feature masking and edge dropout of 0.1 are applied throughout. **Bottom-most – Node features**. Microbe vectors concatenate (i) averaged 2560-d ESM-2 protein embeddings, (ii) 72-d log_1+_ pathway scores, and (iii) 101-d metabolite fingerprints, yielding 2733 features before projection. Metabolite and pathway nodes start from random 128-d and 256-d embeddings, respectively, that are linearly mapped to 768-d.

**Figure 2.**
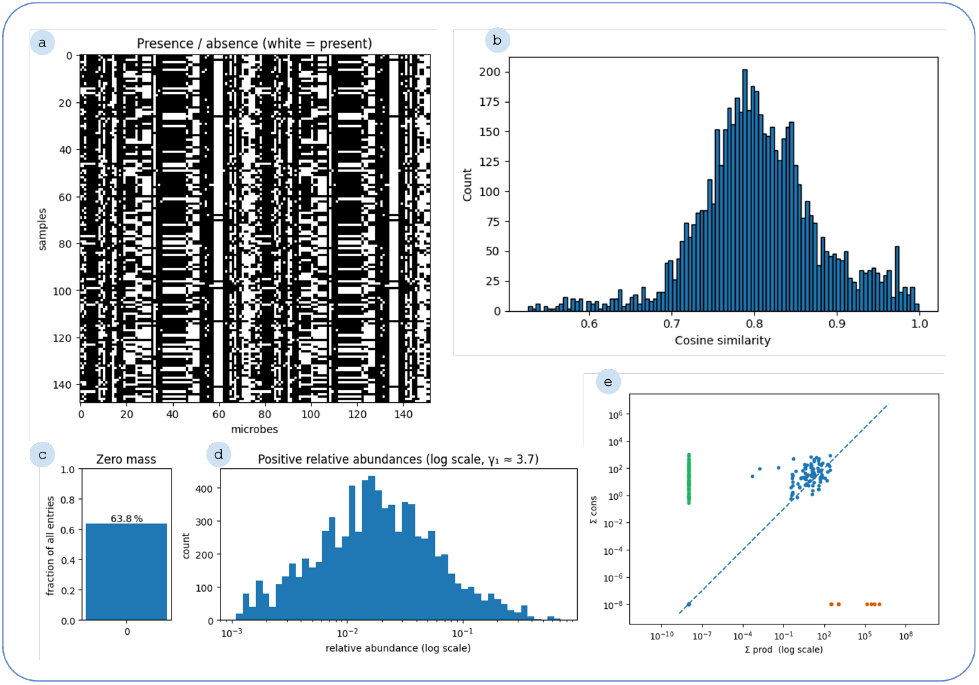
Microbial abundance patterns and metabolic interaction landscape. (a) Presence/absence heatmap of microbes across samples, showing distinct distribution patterns. (b) Histogram of cosine similarities between pathway scores profiles, indicating varying degrees of similarity across samples. (c) Fraction of zero entries (63.8%) in the microbial abundance matrix, illustrating data sparsity. (d) Distribution of nonzero relative abundances on a log scale, showing a heavy-tailed distribution with a shape parameter of 3.7. (e) Comparison of total simulated production versus consumption for each metabolite, which highlights potential metabolic sources and sinks in the microbial community.

#### 2.3.2. Node and edge features

Microbe nodes were characterized using concatenated feature vectors incorporating:

- Protein-level embeddings, generated *via* the ESM-2 model (Lin et al., 2023), yielding 2,560-dimensional vectors from the UHGG database (Almeida et al., 2021), averaged across all proteins in each genome (Elnaggar et al., 2023).
- Pathway fingerprints, log-transformed (log1p) to mitigate skewness.
- Metabolite fingerprints, representing binary indicators of metabolite presence based on flux sampling data.

Pathway and metabolite nodes were initialized with randomly learnable embeddings of dimensions 256 and 128, respectively.

#### 2.3.3. Enhanced Heterogeneous Graph Transformer Architecture

We extended the standard Heterogeneous Graph Transformer (HGT) (Hu et al., 2020) architecture by integrating scalar edge attributes directly into the attention mechanism. This enhancement allowed the attention weights to weigh neighbor contributions by both node features and the strength of interactions derived from metabolite fluxes and microbial abundances. Our resulting model, (`SIMBA`), consists of three transformer layers, each with 768-dimensional hidden states and 12 attention heads—hyperparameters optimized through grid search. To promote training stability and reduce overfitting, we applied residual connections, layer normalization, and a dropout rate of 0.2.

Task-specific output heads were designed to predict for multiple prediction objectives:

- Microbial presence was predicted using sigmoid-activated binary classifier
- Microbial abundances were estimated using a PyTorch Geometric-based softmax layer.
- Metabolite fingerprints were predicted as probabilistic outputs over binary metabolite production capability

The presence head outputs binary labels indicating whether a microbe is expected to occur in a given sample, while the abundance and metabolite heads yield normalized probabilities for species abundances and metabolite production within experimental or simulated communities.

#### 2.3.4. Training Strategy and Loss Functions

We structured our training pipeline into three distinct stages to enhance learning efficacy:

##### Self-supervised learning (SSL)

This initial stage employed graph contrastive learning (GraphCL) (Yang et al., 2024) with a temperature *τ* = 0.10 and a large batch of 256 negatives provides task-agnostic node embeddings.

##### Supervised pretraining

We utilized simulated data to train the model on microbial abundances and metabolite presence predictions. Microbial abundances were optimized using the Tweedie loss with a power parameter of 1.5, selected after comparing results with power values of 1.1, 1.5, and 1.8. The Tweedie loss is defined as:

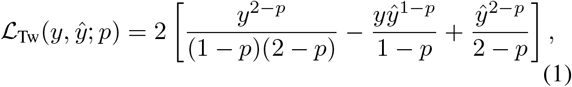

where *y* denotes the true abundance, *ŷ* the predicted abundance, and *p* the power parameter (*p* = 1.5). Metabolite presence (metabolite cross-feeding probability) was optimized with binary cross-entropy (BCE) during supervised pretraining.

##### Fine-tuning

While our model supports joint training on microbial and metabolite outputs, the BSE loss term for metabolite cross-feeding was disabled (*α*_flux_ = 0) for this dataset due to the absence of metabolite-level labels.

We retained the BCE loss for microbial presence prediction and the Tweedie loss for abundance regression. Also, we evaluated a pairwise hinge ranking loss (margin 0.1) with weighting factors *α*_rank_ ∈ {0.05, 0.10, 0.20}. However, using the ranking loss did not yield any improvements in the validation Spearman correlation. As a results, the final model was trained with *α*_rank_ = 0.

Feature masking and edge dropout of 0.1 are applied in every stage.

The stage-specific total loss is

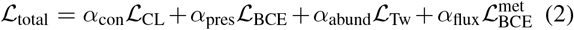

#### 2.3.5. Baseline GNN Models

We compared our enhanced heterogeneous model against baseline homogeneous architectures, including Massage Passign Graph Neural Network (MPGNN), Structured World Models (SWM-GNN) (Kipf et al., 2019), Graph-SAGE (Hamilton et al., 2017), and Simple-GNN. These models utilized only microbial node features and metabolite production fluxes, without explicit edge-value integration. Due to their tendency to overfit, these baseline models were maintained smaller than the heterogeneous model.

#### 2.3.6. Hyperparameter optimization and evaluation metrics

Bayesian hyperparameter optimization was used to identify optimal model configurations. Spearman’s rank correlation served as our primary evaluation metric, as it effectively captures the relative ordering of microbial abundances—an essential aspect of ecological validity. The final model configuration included a hidden dimension of 768, dropout rate of 0.2, edge dropout of 0.1, feature masking rate of 0.1, and 12 attention heads.

Performance evaluation encompassed additional metrics, including presence detection and Spearman score, to ensure comprehensive assessment and biological interpretability. All methods and computational resources are publicly available on GitHub (https://github.com/mofradlab/simba), with complete documentation for reproducibility.

## 3. Results

### 3.1. Characterization of microbial community data and interactions and baseline model performance

To build a comprehensive graphical representation of the gut microbiome for predictive modeling, we first characterized key aspects of our experimental samples and the outputs from pairwise metabolic simulations. These data collectively informed the structure and features of our heterogeneous graph. Figure 2 provides an overview of this foundational data characterization.

The microbial landscape across the samples reveals distinct presence and absence patterns for various microbes (Figure 2a). Understanding these abundance distributions is important as they represent the primary predictive target for our models. In addition, insights into metabolic interplay were derived from pairwise simulations. Figure 2 contrasts the net production and consumption capabilities per metabolite across all microbes in the community which reveals the potential sources and sinks within the simulated ecosystems and underpins the microbe-metabolite interactions edges in our graph.

Collectively, these analyses underscore the heterogeneity of microbial membership, the complexity of functional similarities, the challenging nature of abundance distributions, and the intricate web of simulated metabolic interactions. This rich, multi-modal information landscape motivated the use of GNNs capable of integrating diverse data types to model microbial community behavior.

To establish a performance benchmark, we initially evaluated several established GNN architectures: SWM-GNN, GraphSAGE, MPGNN, and GNN. As illustrated in Figure S3, the models showed modest performance in predicting microbial abundances, with Spearman rank correlations generally below 0.6. This highlighted the limitations of standard GNNs for this complex task and motivated the development of our specialized `SIMBA` architecture.

### 3.2. Performance across the training pipeline

To address the challenges identified, we propose `SIMBA` to predict microbial community composition. This model is developed to predict microbial presence and relative abundances by leveraging information from simulated metabolic cross-feeding interactions within an HGT framework. The model’s performance was established through a three-stage training regimen (self-supervised learning, supervised pretraining, and fine-tuning) and its architecture was refined *via* systematic hyperparameter optimizatrion.

#### 3.2.1. Pretraining on simulated data

Following SSL, `SIMBA` was pretrained on a comprehensive dataset of simulated pairwise microbial interactions, enablinging the model to learn the fundamental patterns of microbial metabolic cross-feeding. The convergence of the training loss over epochs is shown in Figure S4. The total training loss, alongside its components, BCE loss, Tweedie loss, and rank loss, all showed a consistent decrease over epochs of pretraining, indicating stable learning dynamics. For the auxiliary task of predicting metabolite cross-feeding presence, the model attained an accuracy of 0.96. The corresponding F1-score and recall for this task were 0.83 and 0.72, respectively, which further supports the model’s capability to identify these interactions in the simulated data.

Collectively, the results from the supervised pretraining stage indicate that SIMBA successfully learned to model key aspects of microbial ecology within the context of the simulated data. This provided a strong foundation for the subsequent fine-tuning stage on experimental data.

#### 3.2.2. Fine-tuning on experimental data

The final stage involved fine-tuning the pretrained `SIMBA` model using experimental microbial abundance data. This allowed the model to adapt its learned representations and predict capabilities to real-world community contexts. The fine-tuning loss progression is presented in Figure S5. Critically, the Spearman correlation of microbial abundance prediction on the experimental validation set showed a consistent improvement over fine-tuning epochs, reaching 0.85. This highlights our model’s ability to effectively transfer knowledge from simulated environments and adapt to experimental data complexities.

### 3.3. Development and optimization

The optimal configuration was obtained using Bayesian hyperparameter optimization, with the primary objective of maximizing the Spearman rank correlation for microbial abundance prediction. This process was done by tuning key hyperparameters including model’s hidden dimension, attention head, edge dropout, feature masking, the weight of the rank loss component in the combined loss function. The overall relationship between the hyperparameters and the achieved Spearman correlation is shown in Figure S6. Specifically, varying the feature masking revealed an optimal range around 0.1, where higher masking led to a slight decline in performance. Similarly, an edge dropout centered around 0.2 was found to be most effective for microbial abundance prediction. The influence of the rank loss weight showed stable performance around zero.

Further investigation into play between hidden dimension (`D256`, `D512`, `D768`, and `D1024`) and attention head (`H8`, `H12`, and `H16`) was performed, and `SIMBA-D768-H12` outperformed others with a Spearman correlation above 0.8 (see Figure S7). The Spearman scores for all models showed a trend of stabilizing after approximately 20 epochs. The complete set of optimized hyperparameters employed for `SIMBA` is detailed in Table 2.

**Table 1.** Loss-weight schedule used in the final model.

| Stage | $\alpha_{\text{con}}$ | $\alpha_{\text{pres}}$ | $\alpha_{\text{abund}}$ | $\alpha_{\text{flux}}$ |
| --- | --- | --- | --- | --- |
| Self-supervised (SSL) | 1.0 | 0 | 0 | 0 |
| Supervised pretraining | — | 1.0 | 1.0 | 1.0 |
| Fine-tuning | — | 1.0 | 1.0 | 0 |

**Table 2.**
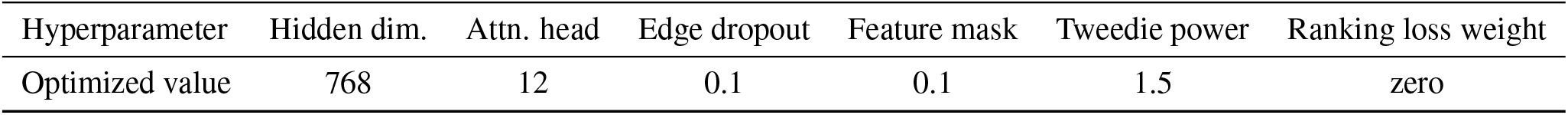
Final optimized hyperparameter configurations for SIMBA. Values were determined through Bayesian hyperparameter optimization targeting Spearman correlation for microbial abundance.

A critical aspect of modeling microbial abundances is the choice of an appropriate loss function, given the characteristic zero-inflation and skewed distribution of such data. We systematically evaluated several loss functions for the abundance prediction task during the fine-tuning stage. From Figure S8 that represents the Spearman correlation achieved with different loss functions, the Tweedie loss with a power parameter of 1.5 demonstrated superior performance in capturing the abundance distribution, outperforming both KL divergence and Huber loss functions.

### 3.4. Microbial and metabolite prediction insights

At an individual level, Figure 3 presents a detailed comparison of ground truth data and model predictions for a representative sample from the experimental samples. The model performs well in predicting both the presence of microbes in the community and the metabolites cross-feeding among them. Absence is indicated by gray circles, while colored circles denote presence. Color intensity in the prediction circles reflects the concordance with ground truth abundance, calculated as a ratio, where higher intensity signifies a closer match. This visualization highlights SIMBA’s ability to accurately identify present microbes and its general agreement with actual abundance levels for many species in this example. Also, the figure provides insight into the model’s predictions related to metabolite cross-feeding probabilities.

**Figure 3.**
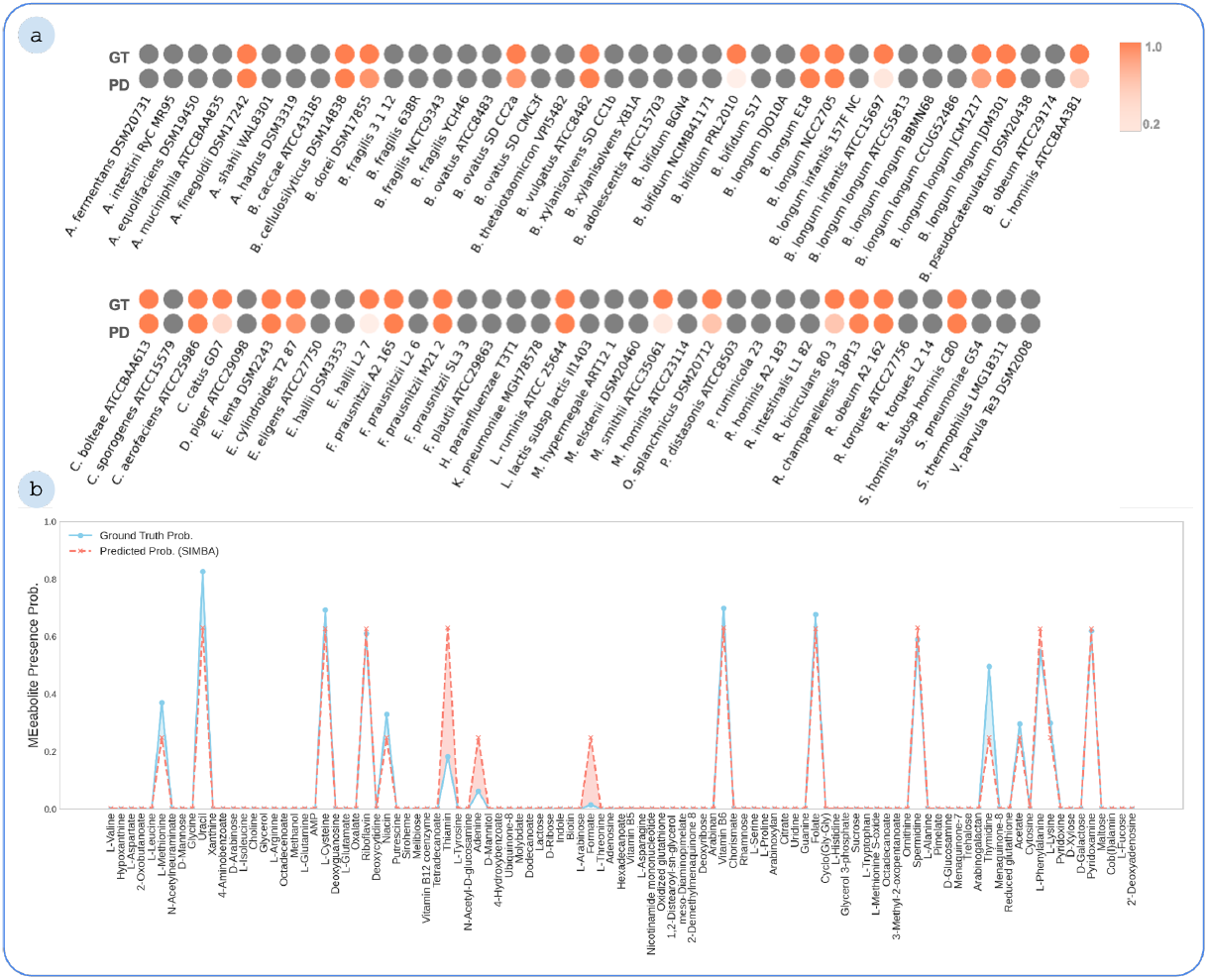
Comparison of ground truth and predictions for an exemplary sample; (a) `SIMBA` performance in predicting microbial presence and abundance. Each pair of circles represents a microbe: the top (GT) shows ground truth and the bottom (PD) shows model prediction. Gray circles indicate absence, while colored circles indicate presence. The color intensity reflects how well the predicted abundance matches the ground truth, computed as a ratio between predicted and actual values. A higher intensity suggests a closer match. (b) Comparison of ground truth probabilities and predicted probabilities for different metabolites. (GD: ground truth; PD: prediction)

## 4. Discussion

Modeling the human gut microbiome is particularly challenging due to the heterogeneous, zero-inflated, and compositionally biased nature of microbial abundance data. Microbial communities are governed not only by taxonomic presence but also by metabolic behavior and ecological interactions, such as cross-feeding, competition, and syntrophy. Traditional modeling approaches that rely solely on abundance data or simplified interaction assumptions fail to capture the underlying microbiological and ecological richness.

To address these challenges, we combined experimental data with mechanistic modeling and graph-based learning to generate a multimodal representation of microbial communities. By simulating 2,850 pairwise interactions using metabolic networks, we captured a rich landscape of potential metabolic cross-feeding—a crucial ecological signal often inaccessible in experimental settings. These simulated interactions were then used to construct a heterogeneous interaction graph, integrating functional embeddings, pathway annotations, and probabilistic metabolite cross-feeding data.

Against this complex data backdrop, we developed a simulation-augmented GNN framework tailored to the microbiome domain. Unlike standard models, which struggled to exceed a Spearman correlation of 0.6, our approach integrates domain-specific knowledge through a heterogeneous graph transformer, modified to incorporate edge attributes directly into the attention mechanism. This enhancement enables biologically grounded modeling of both the strength and directionality of simulated microbial interactions.

The training pipeline mirrors this complexity. By employing a three-stage strategy—self-supervised representation learning, supervised pretraining on simulated data, and fine-tuning on experimental abundance profiles—we facilitated effective knowledge transfer between *in silico* simulations and *in vivo* observations. This progression stabilized learning dynamics and yielded substantial performance gains, ultimately achieving a Spearman correlation of 0.85 on experimental validation data.

Several architectural and training decisions contributed to this success. Bayesian hyperparameter optimization identified an optimal configuration: moderately large hidden dimensions (768), 12 attention heads, and low edge dropout and feature masking rates (both 0.1). The Tweedie loss (power = 1.5) was especially effective in modeling the zeroinflated, skewed abundance distributions, outperforming alternatives like KL divergence and Huber loss.

Beyond predictive accuracy, the framework also supports microbiological interpretability. By combining species presence with their metabolic capacities and cross-feeding patterns, it enables identification of keystone taxa and metabolic bottlenecks—offering a mechanistic foundation for developing targeted therapeutic or dietary interventions.

## 5. Conclusion

Our results demonstrate that a biologically informed, simulation-augmented GNN architecture can substantially improve microbial abundance prediction. SIMBA represents a step toward ecologically grounded, mechanistically interpretable models of microbial communities. Using domain knowledge, metabolic simulations, and advanced graph learning, SIMBA offers a robust tool for both microbiome prediction and hypothesis generation in microbial ecology.

### Impact Statement

This paper presents SIMBA, a novel framework for predicting gut microbial interactions and community composition. The primary goal is to advance our understanding of complex microbial ecosystems and their role in health and disease. Potential societal benefits include improved personalized nutrition and therapeutic strategies for microbiome-related conditions. In this work, we have focused on mechanistic interpretability and robust evaluation.

## A. Appendix

**Figure S1.**
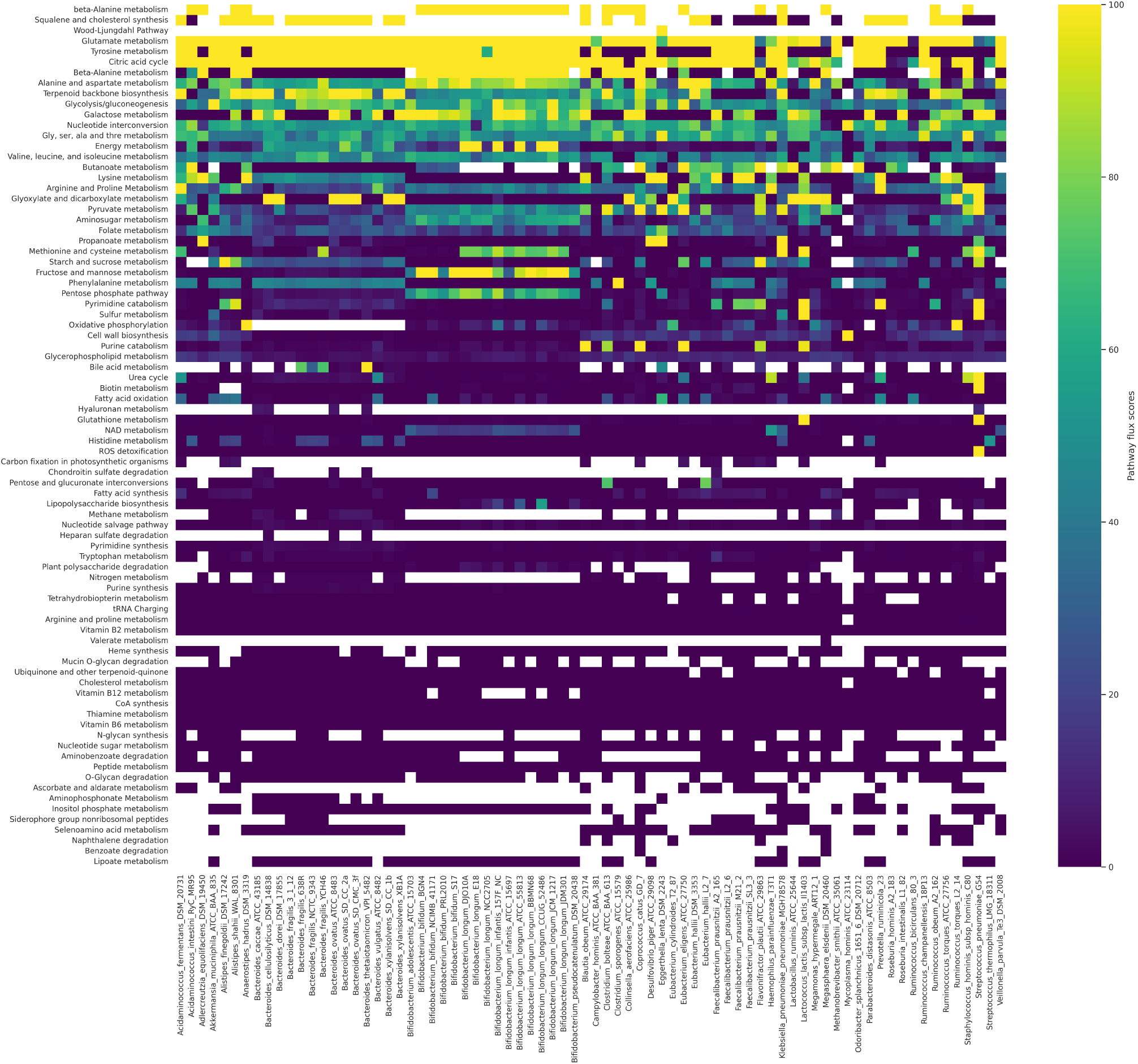
Heat map representation of pathway scores for microbes involved in simulations. Rows represent metabolic pathways, columns represent microbial strains, and cell intensity indicates pathway activity scores.

**Figure S2.**
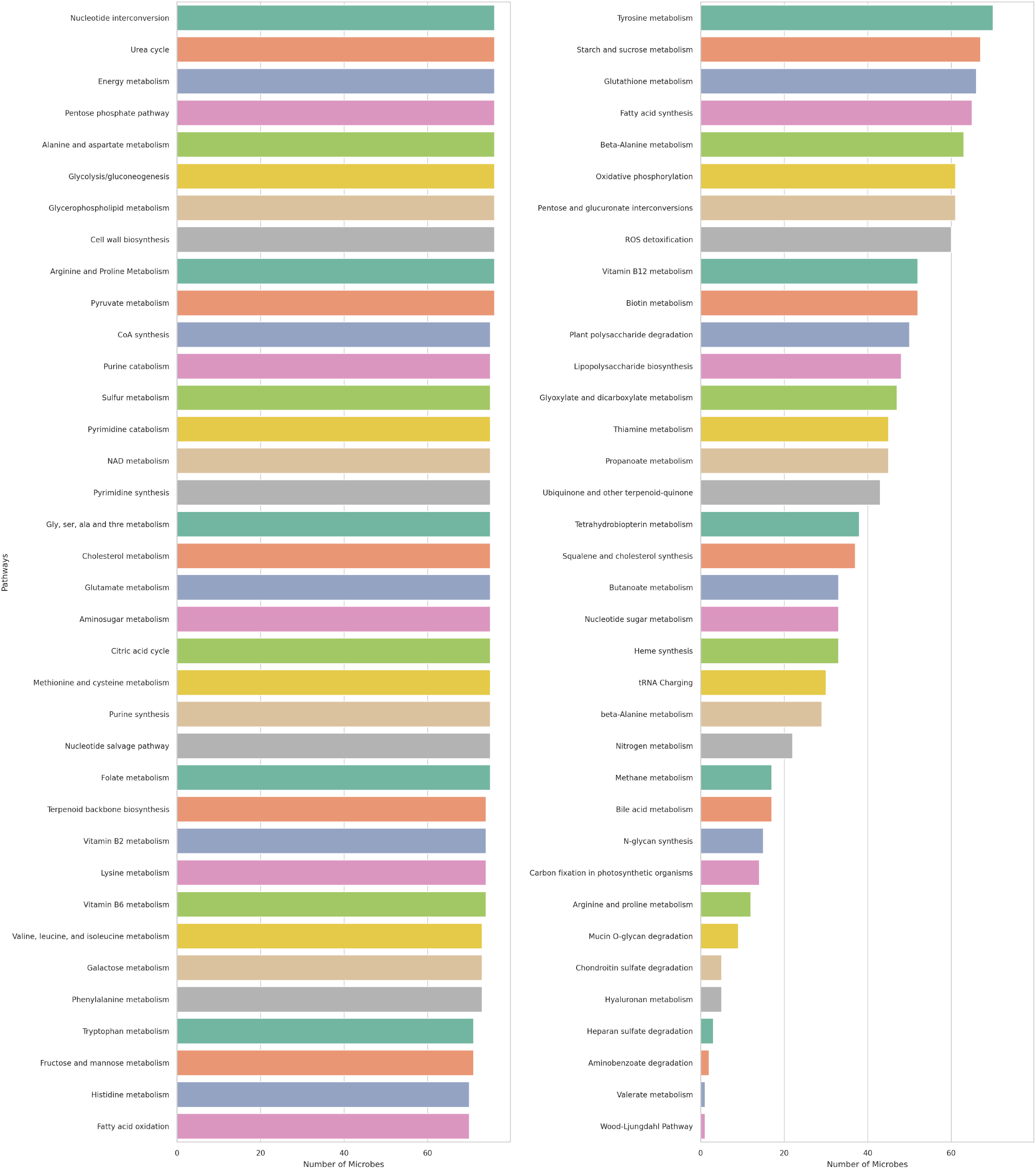
Pathway distributions for strains obtained from pairwise simulations. Bar charts showing the number of microbes associated with various metabolic pathways.

**Figure S3.**
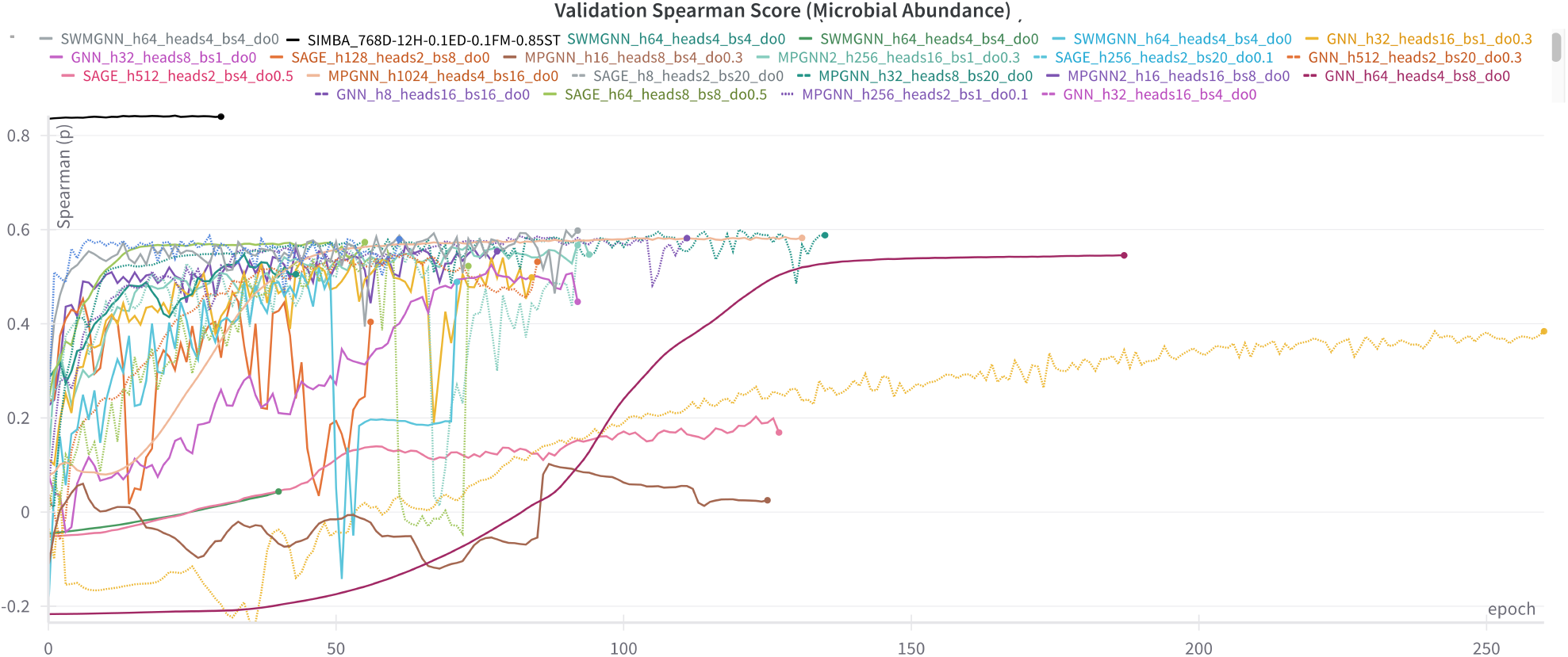
Different GNN baseline model Spearman scores. The models show limited performance in microbial abundance prediction.

**Figure S4.**
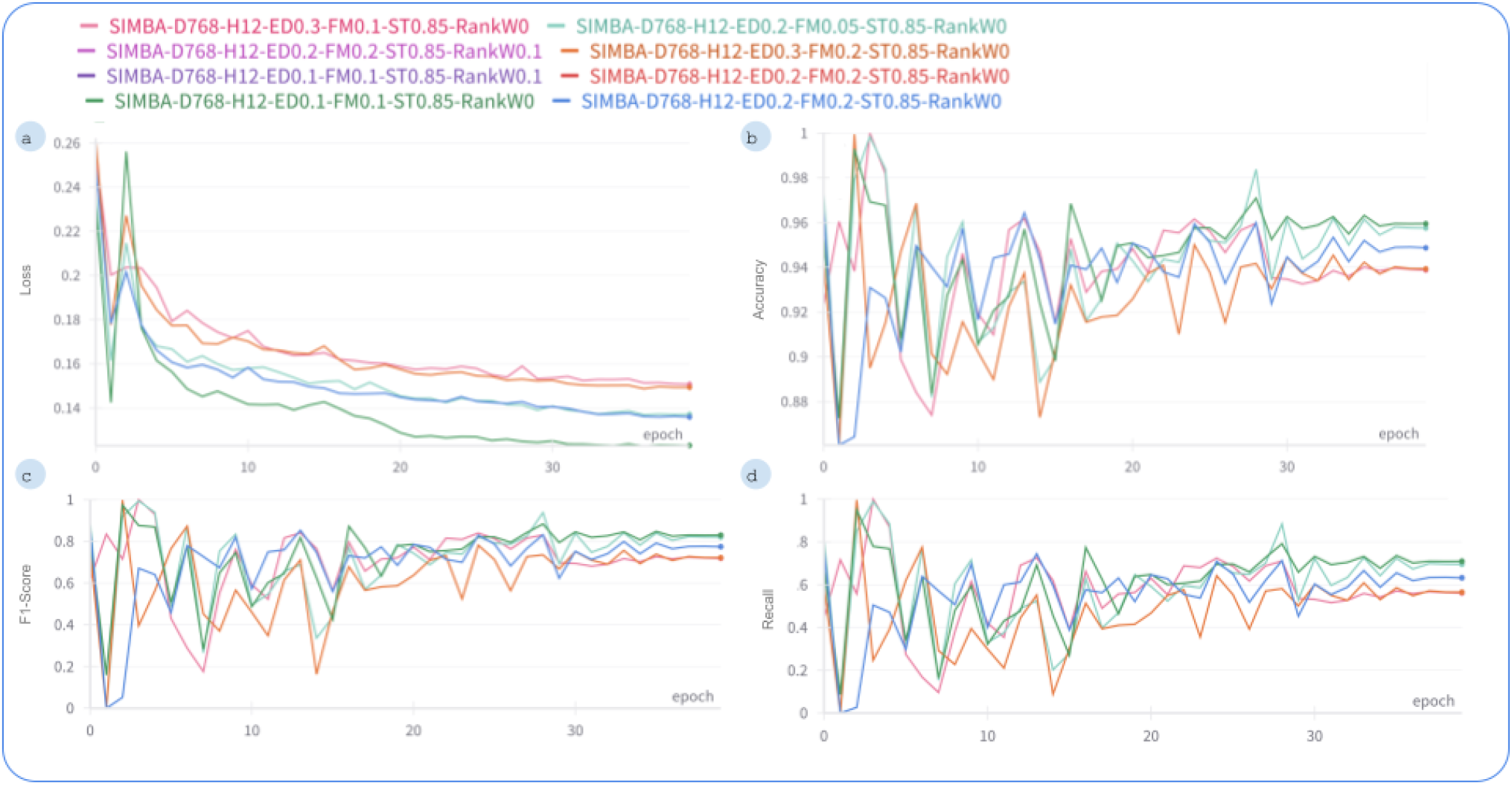
Performance of `SIMBA` during supervised pretraining on simulated data. (a) Convergence of training loss over epochs. (b-d) Performance on the validation set for predicting the presence of metabolite cross-feeding presence over epochs, showing (b) accuracy, (c) F1-score, and (d) recall.

**Figure S5.**
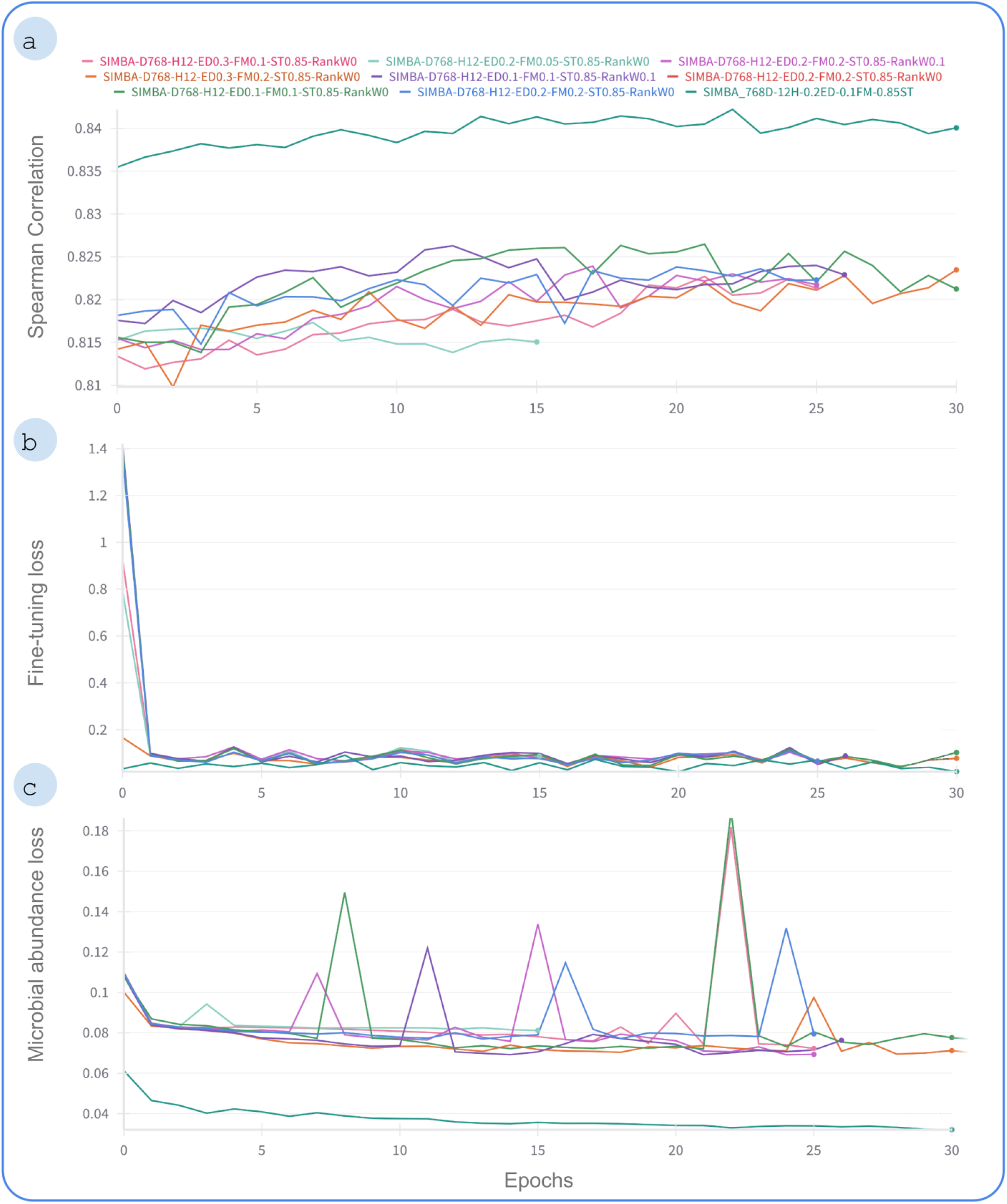
Performance `SIMBA` during fine-tuning on experimental data. (a) Fine-tuning validation performance of the model variants for microbial abundance prediction. Spearman correlation trajectories on the experimental validation set are shown over the course of fine-tuning epochs. (b) Convergence of fine-tuning loss over epochs. (c) Convergence of microbial abundance loss during fine-tuning.

**Figure S6.**
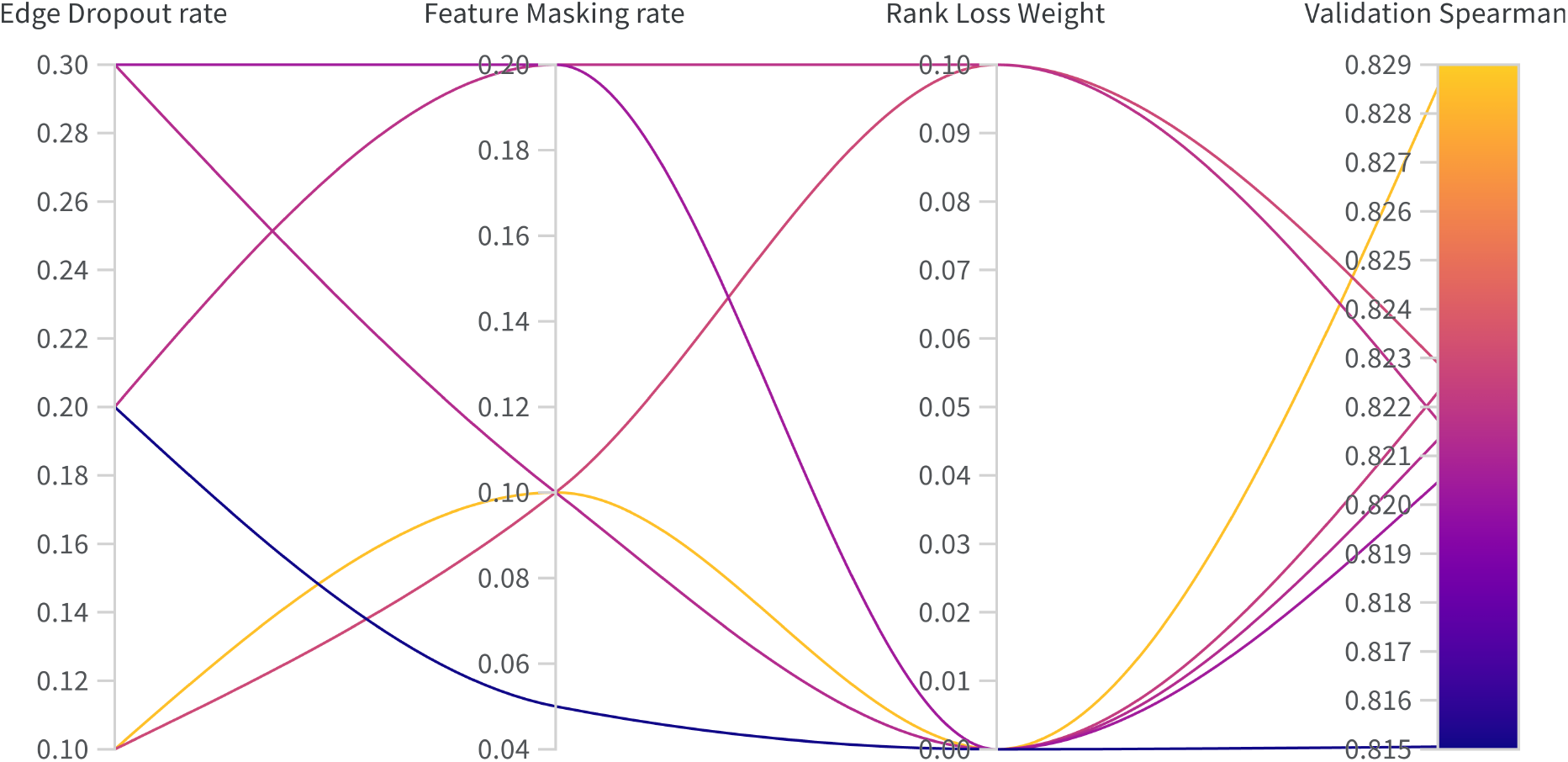
Relationship between key hyperparameters and the achieved Spearman scores for microbial abundance on the validation set during Bayesian hyperparameter optimization.

**Figure S7.**
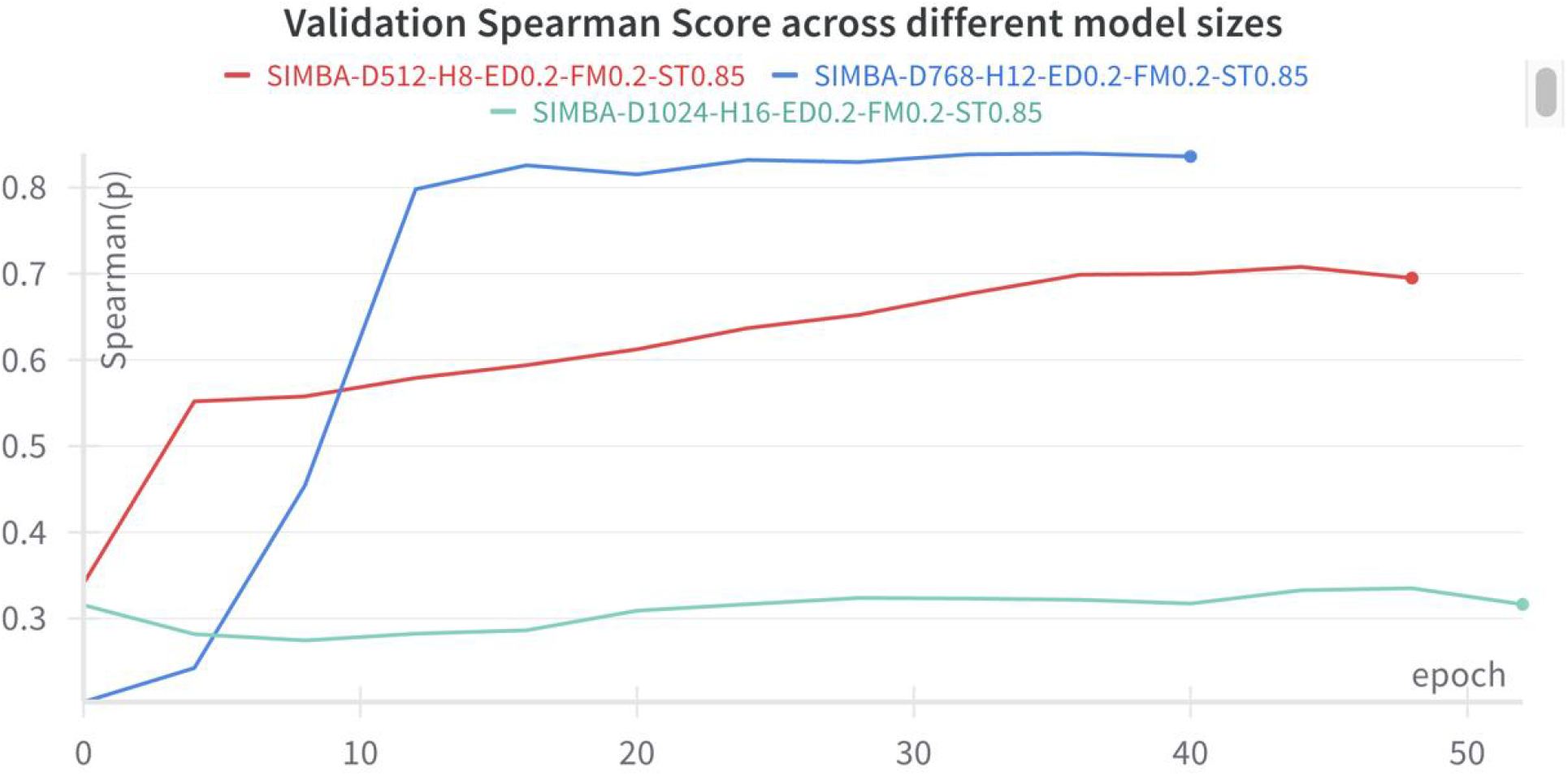
Changes of Spearman scores for models with different hidden dimensions and attention heads.

**Figure S8.**
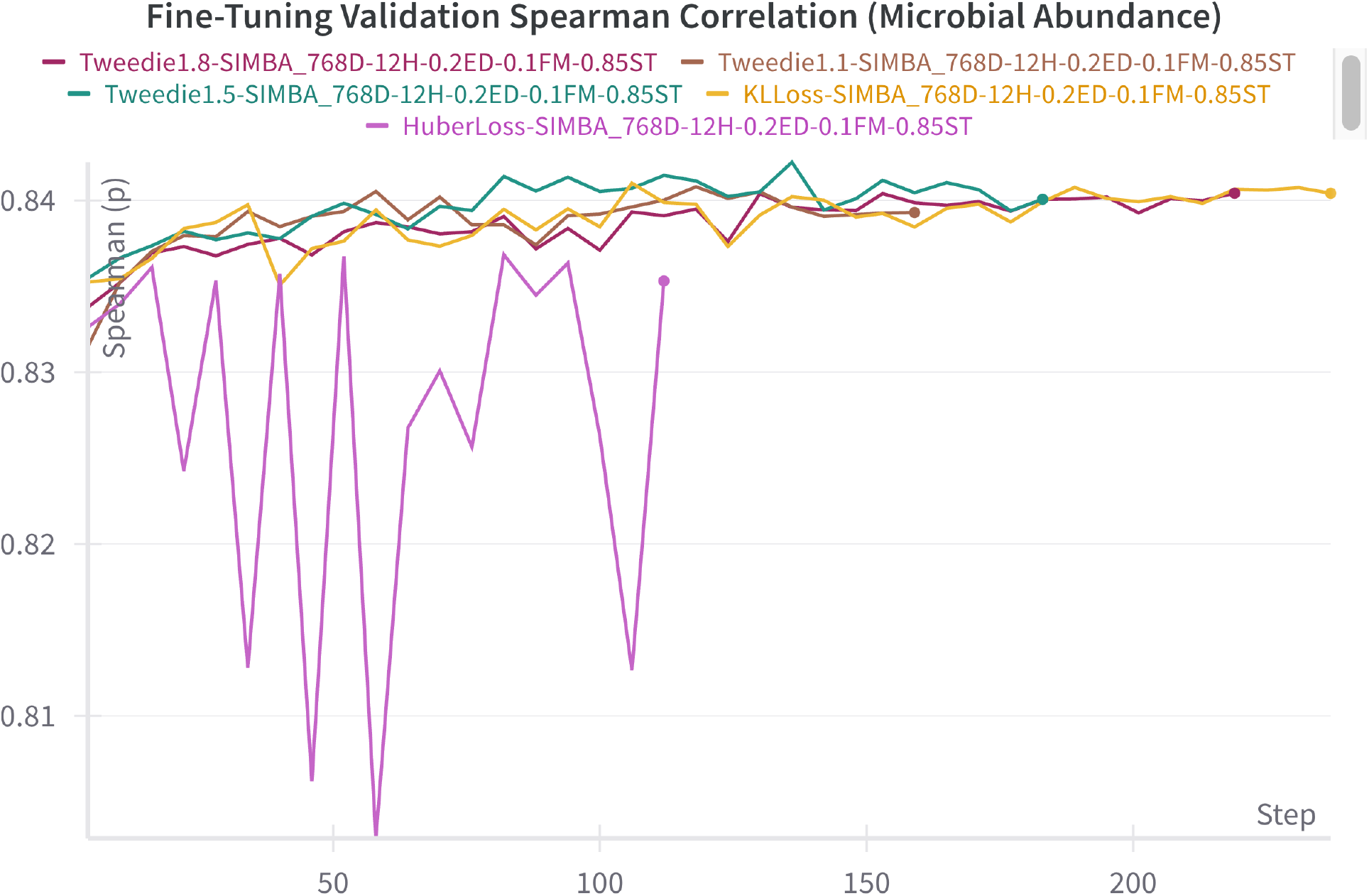
Comparison study of different loss functions used for the prediction of microbial abundances based on Spearman correlation

